# SNAP-tagged nanobodies enable reversible optical control of a G protein-coupled receptor via a remotely tethered photoswitchable ligand

**DOI:** 10.1101/266247

**Authors:** Helen Farrants, Amanda Acosta Ruiz, Vanessa A. Gutzeit, Dirk Trauner, Kai Johnsson, Joshua Levitz, Johannes Broichhagen

## Abstract

G protein-coupled receptors (GPCRs) mediate the transduction of extracellular signals into complex intracellular responses. Despite their ubiquitous roles in physiological processes and as drug targets for a wide range of disorders, the precise mechanisms of GPCR function at the molecular, cellular, and systems levels remain partially understood. In order to dissect the function of individual receptors subtypes with high spatiotemporal precision, various optogenetic and photopharmacological approaches have been reported that use the power of light for receptor activation and deactivation. Here, we introduce a novel and, to date, most remote way of applying photoswitchable orthogonally remotely-tethered ligands (PORTLs) by using a SNAP-tag fused nanobody. Our nanobody-photoswitch conjugates (NPCs) can be used to target a GFP-fused metabotropic glutamate receptor by either gene-free application of purified complexes or co-expression of genetically encoded nanobodies to yield robust, reversible control of agonist binding and subsequent downstream activation. By harboring and combining the selectivity and flexibility of both nanobodies and self-labelling enzymes, we set the stage for targeting endogenous receptors *in vivo*.

## Introduction

The optical control of proteins and their associated cellular functions has evolved into a powerful technique for gaining a deeper understanding of biological processes[1, 2]. In particular, the ability to optically control ion channels, transporters and receptors has made it possible to move toward a more mechanistic understanding of the role of specific cells and synapses in the circuit-driven processes of the brain. While classical optogenetic tools, such as those based on opsins, allow for the general control of cellular excitability or broadly-defined signaling pathways, the ability to control specific signaling molecules has the potential to extend such analysis to probe the *molecular* basis of physiological functions and disease pathophysiology. This circumvents the limitations of pharmacology, where subtype-selectivity is difficult to achieve and compounds cannot be applied or removed in a fast and efficient manner to defined locations due to diffusion.

Ideally, one would be able to control *native* receptors with subtype-specificity, high spatiotemporal precision, and genetic targeting to defined cell types. A number of previous strategies have achieved a subset of these desired parameters, but it remains a major challenge to attain all of them within the same toolset[3] (**Supplementary Table 1**). We previously developed approaches to optogenetically manipulate specific G protein-coupled metabotropic glutamate receptors (mGluRs) through the covalent attachment of photoswitchable tethered ligands (PTLs)[4-6]. Initially, these compounds were targeted directly to an engineered cysteine *via* maleimide conjugation (Fig 1A)[7], but more recently we improved this method through the use of an *N*-terminal self-labeling tag (Fig 1B)[8]. This strategy has yielded enhanced labeling efficiency in culture and *in vivo*[9], simplified photoswitch design, enabled multiplexing of tags, and, most importantly, is fully orthogonal to native chemistry[10]. Based on the demonstrated utility of self-labeling proteins in this context and the key finding that tethered photoswitches can optically control ligand binding when attached to a distinct protein domain, we wondered if we could further transport photoswitch attachment to an antibody or antibody fragment to add an extra layer of specificity and flexibility (Fig 1C). The ability to photoswitch a membrane receptor in this manner would produce an approach that could be used to target native receptors without the need for the incorporation of a labeling tag directly into the protein of interest. To achieve this, we turned to single chain antibodies (*i.e.* nanobodies). Nanobodies are beneficial because of their small size, target specificity, and ability to be genetically encoded for expression in mammalian cells[11, 12].

**Fig 1.**
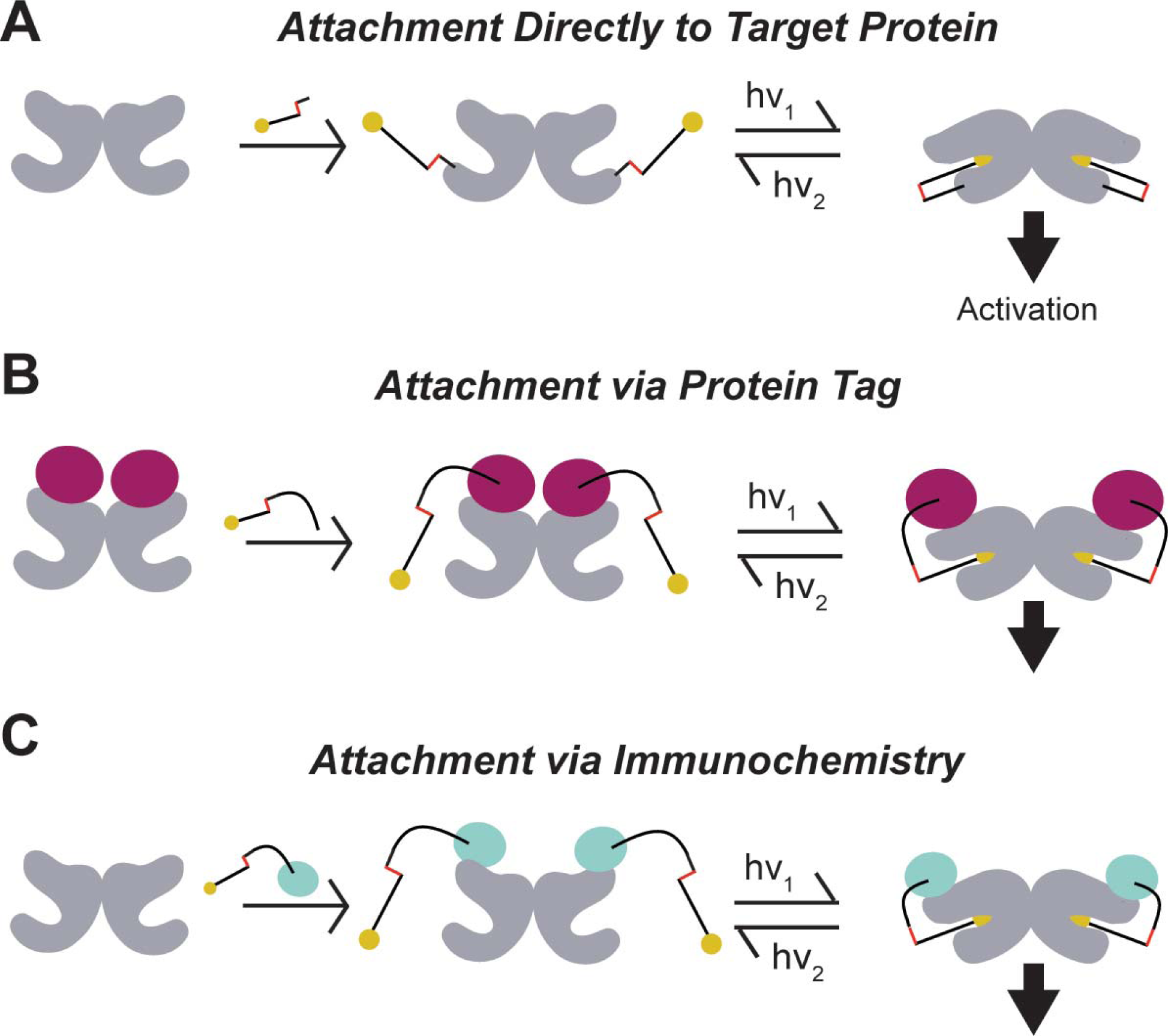
Optical control of ligand binding via distinct photoswitch attachment strategies. Photoswitchable tethered ligands may be presented to target ligand binding domains through previously established strategies of directly binding to the target protein (A), traditionally *via* cysteine-maleimide linkage, through binding to introduced self-labeling protein tags (B), such as SNAP, CLIP or Halo-tag, or potentially through immunochemistry with labeled antibodies (C).

## Results

### SNAP-tagged anti-GFP nanobodies (NBs) retain GFP affinity and efficient labeling of BG-conjugated compounds

To enable gene-free optical control of target proteins, we sought to utilize previously characterized nanobodies (NBs) that target GFP[13] to produce a “Nanobody-Photoswitch Conjugate” (NPC; see **Supplementary Table 1**). We genetically fused the self-labeling SNAP-tag[14, 15] protein to either the *N*-terminus (“SNAP-NB”) or *C*-terminus (“NB-SNAP”) of the anti-GFP nanobody GBP1, which enhances the fluorescence of wtGFP. The SNAP-fusion proteins remained reactive in this context and allowed us to covalently attach small molecules, such as fluorophores and photoswitches, to the NBs *via O*^6^-benzylguanine (BG)-linked compounds *in vitro* (**Fig S1**). We next tested if SNAP incorporation would alter the ability of the NB to efficiently target its antigen, GFP. Application of both SNAP-NB and NB-SNAP fusions to purified wtGFP led to a dose-dependent increase in fluorescence (Fig 2A), with an apparent affinity of ∼3 nM. Together this shows that incorporation of SNAP into the NB at either end does not affect its ability to target GFP, consistent with the localization of the *N* and *C*-termini of the NB distant from the GFP binding site[13].

**Fig 2.**
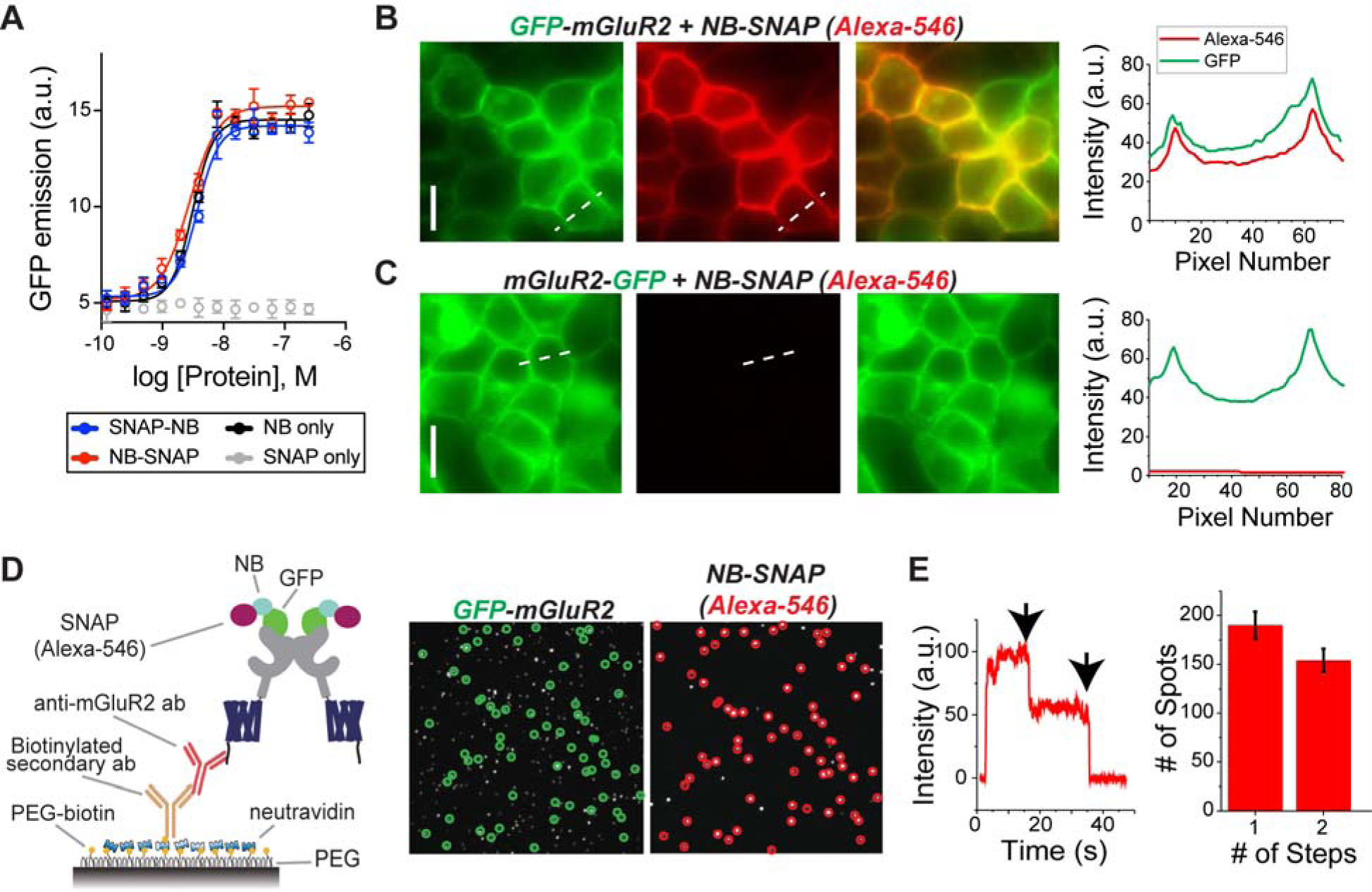
SNAP-tagged anti-GFP nanobodies enable stoichiometric labeling of GFP-mGluR2 in living cells. (A) *In vitro* dose-response curve for SNAP-NB, NB-SNAP, NB and SNAP against wtGFP assessed by fluorescence enhancement upon binding indicates that attachment of *N* or *C*-terminal SNAP domains does not alter NB affinity (error bars indicate SEM). (B) Representative images showing labeling of surface-expressed GFP-mGluR2 (green) by application of purified NB-SNAP labeled with Alexa-546 (red). Right, line scan analysis shows overlap in GFP and Alexa-546 fluorescence at the plasma membrane. (C) Representative images, left, and line scan, right, showing that no red surface fluorescence is observed following application of Alexa-546-labeled NB-SNAP to cells expressing mGluR2-GFP. (D) Left, Schematic of single molecule pulldown (SiMPull) of GFP-mGluR2 and SNAP-fused NBs *via* an anti-mGluR2 antibody on polyethylene-glycol (PEG) passivated coverslip. Right, representative images show co-localized GFP-mGluR2 (green) and NB-SNAP (Alexa-546; red) molecules. 74.6% of red spots were colocalized with a green spot (773/1034). (E) Counting photobleaching steps in the red channel (representative trace, left) allow for analysis of NB stoichiometry (bar graph, right), confirming that 2 NBs can bind to a receptor dimer.

### SNAP-tagged NBs label GFP-mGluR2 in living cells with 2:2 stoichiometry

We next asked if purified SNAP-tagged NBs could enable efficient labeling of a GFP-tagged membrane receptor to ultimately mediate optical control in living cells. We genetically fused GFP to either the extracellular *N*-terminus or the intracellular *C*-terminus of mGluR2 (see online methods). Application of purified SNAP-tagged NBs that had been pre-labeled with Alexa-546 produced clear fluorescence on the surface of HEK 293T cells expressing GFP-mGluR2, but not mGluR2-GFP (Fig 2B,C; Fig S2A,B). Importantly, following cell surface labeling with NBs, washing with buffer for 30 minutes did not reduce the fluorescence (**Fig S2C**), indicating that the extracellular GFP-NB interaction is sufficiently stable for probing signaling on the physiological timescales relevant to GPCR signaling.

Many membrane receptors are oligomeric, raising the need for labeling of each subunit within a complex if one is going to use NBs for efficient optical control of ligand binding with a tethered agonist. Since mGluRs are dimers in living cells[16, 17] and two agonists are required for full activation[16, 18], we sought to determine if two NBs can bind to an mGluR2 dimer. Compared to typical antibodies (∼150 kDa), we hypothesized that the small size of NBs (∼15 kDa) would decrease the chance of steric hindrance between NBs on adjacent subunits. To measure the stoichiometry of SNAP-tagged NBs bound to GFP-mGluR2 dimers, we performed a single-molecule pulldown (“SiMPull”) of lysate from cells expressing GFP-mGluR2 and labeled with NB-SNAP(Alexa-546). After immobilization of GFP-mGluR2 on a polyethylene-glycol (PEG) passivated coverslip using an anti-mGluR2 antibody (Fig 2D), we used photobleaching step analysis of spots in the red channel that were co-localized with GFP spots to determine the number of NBs per GFP-mGluR2 (Fig 2D,E). ∼45% of these colocalized red spots bleached in 2-steps, showing that two NBs can indeed bind to an mGluR2 dimer (Fig 2E). Importantly, very few spots were observed when lysate from cells expressing mGluR2-GFP was immobilized (**Fig S2D**).

### Purified SNAP-tagged NBs allow photoactivation of mGluR2 via “BGAG” photoswitches

Given the efficient binding of SNAP-tagged NBs to GFP-mGluR2, we next tested if they could enable optical control of mGluR2 *via* attachment of BGAG photoswitches[8, 10]. As a pre-requisite to optical control, we first assessed the ability of GFP-mGluR2 to retain normal function following NB conjugation. We performed whole cell patch clamp recordings from HEK 293T cells co-expressing G protein-gated GIRK channels as a readout of activity and saw no effect of NB attachment on apparent glutamate affinity or maximum current amplitude (Fig 3A). We next labeled GFP-mGluR2 expressing cells with either SNAP-NB or NB-SNAP along with BGAG variants of different PEG linker length (BGAG_0_, BGAG_12_, or BGAG_28_) (Fig. 3B). Robust photocurrents were observed for most conditions with maximal photo-activation of up to 40% relative to saturating glutamate observed for NB-SNAP with BGAG_12_ (Fig 3C; S3A-C). The directionality of photoswitching, with *cis*-BGAG serving as an agonist, was conserved for NB conjugation to all BGAG variants as was seen in previous studies with direct conjugation to an *N*-terminal SNAP domain[8, 10]. This supports a mechanism where relative efficacy of the *cis* and *trans* configurations are due to alterations in the inherent pharmacology of the azobenzene-glutamate moiety, rather than the relative reach of the two forms. Importantly, in the presence of saturating (1 mM) glutamate, no photoswitching was observed indicating that BGAG_12_ does not serve as a partial agonist or antagonist in the NB-tethered context (**Fig S3D**). Subtle differences in the BGAG length dependence of SNAP-NB and NB-SNAP indicate differences in the relative orientation to the glutamate binding site. SNAP-NB showed similar efficacy with BGAG_0_ and BGAG_12_ whereas NB-SNAP showed a 2-fold increase in efficacy for BGAG_12_ relative to BGAG_0_ (Fig 3D). This likely indicates that the SNAP domain of SNAP-NB is, on average, closer to the glutamate binding site than the SNAP domain of NB-SNAP. Both SNAP-NB and NB-SNAP showed negligible photoswitching with BGAG_28_, consistent with the notion that PEG chains of this length provide too low of an effective concentration due to the large volume of conformational space sampled. Structural models showing a possible arrangement of SNAP-tagged nanobodies relative to GFP-mGluR2 indicate distinct geometries of the two systems which could account for differences in photoswitch efficacy (**Fig S3E**). Protein flexibility likely also plays a major role in facilitating the ability of NB-tethered BGAGs to effectively bind and activate mGluR2. This is especially likely for BGAG_0_ which would be too short to reach the binding site without major flexibility introduced either from inter-domain linkers (*i.e.* between GFP and mGluR2 or SNAP and NB) or the receptor itself.

**Fig 3.**
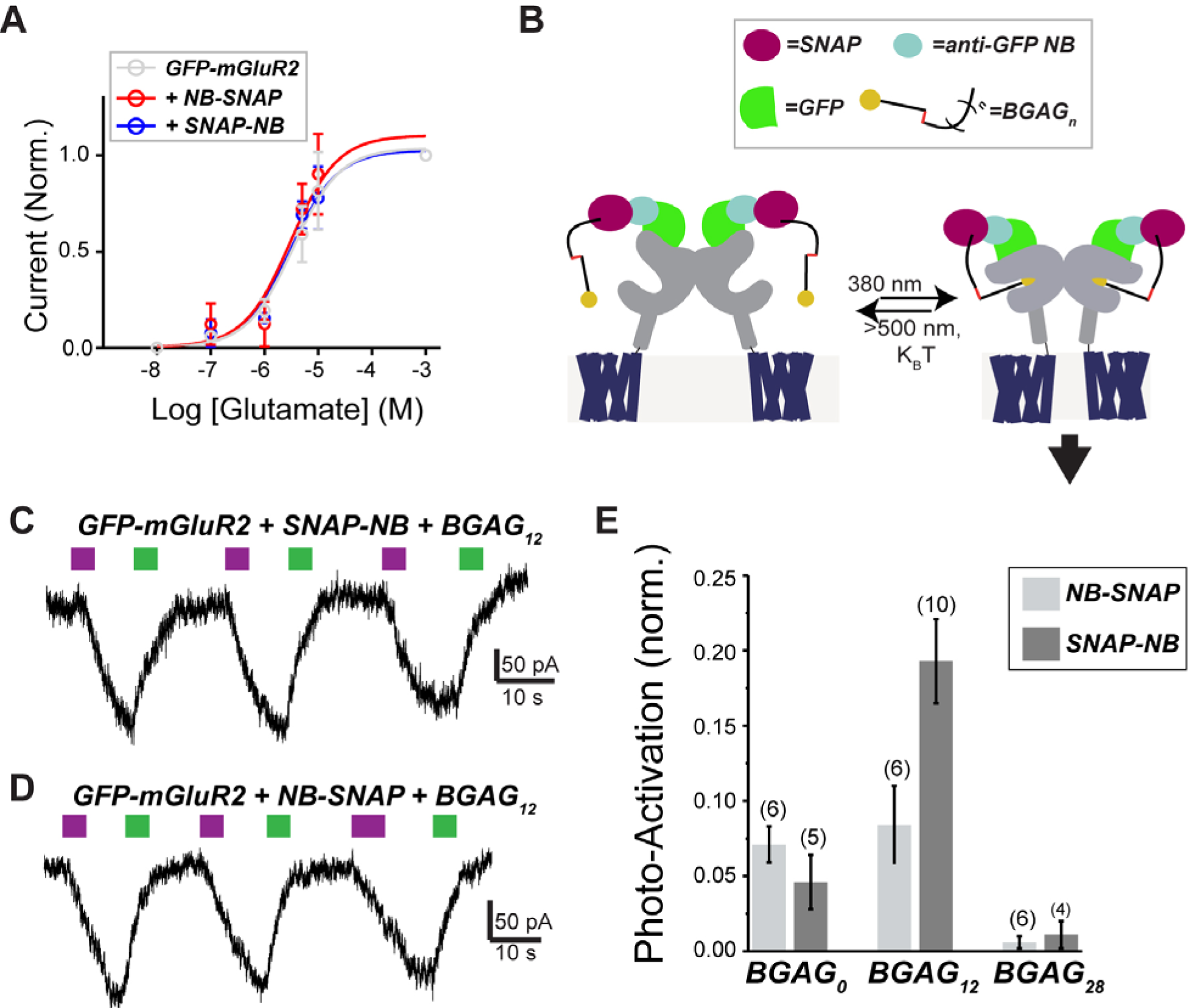
BGAG conjugation to SNAP-tagged NBs permits robust photo-activation of GFP-mGluR2. (A) Glutamate dose-response curves, from GIRK activation assay, indicate that conjugation of SNAP-NB or NB-SNAP does not change the glutamate response of mGluR2. Current amplitudes are normalized to the response to saturating 1 mM glutamate. (B) Schematic showing arrangement of BGAG-labeled SNAP-tagged NBs attached to GFP-mGluR2 and light-mediated receptor activation via photoisomerization of the azobenzene moiety of BGAG. (C) Representative traces showing reversible photoactivation of GFP-mGluR2 by BGAG_12_ conjugated to SNAP-NB (top) or NB-SNAP (bottom). 380 nm light (magenta) induces photo-isomerization of the azobenzene from *trans* to *cis* and 500 nm light (green) reverses the process. (D) Summary bar graph showing relative efficacy for BGAG variants of different length conjugated to either SNAP-NB or NB-SNAP. The numbers of cells tested for each condition are shown in parentheses.

### Co-expression of genetically encoded SNAP-tagged NBs enables optical control of mGluR2

While the ability to target proteins for optical control using purified nanobodies opens many experimental possibilities that don’t require genetic manipulation, an alternative approach would take advantage of the relative ease at which NBs can be genetically encoded and expressed in mammalian systems[19, 20]. Genetically encoding NBs that can then target specific proteins and provide a tethering point for photoswitch attachment would allow for cell-type specific targeting, a crucial property for dissecting the role of specific signaling molecules in many physiological systems, especially within the circuitry of the nervous system. To see if NBs can be used to target extracellular protein domains, we added an mGluR signal sequence (ss) to NB-SNAP to promote trafficking through the endoplasmic reticulum (ER) *via* the same pathway as integral membrane proteins. This should allow NB-SNAP to remain in the same compartment (ER lumen, Golgi lumen, extracellular space) as the GFP of GFP-mGluR2 during the entire trafficking process (Fig 4A). Consistent with this, co-expression of ss-NB-SNAP and GFP-mGluR2, but not mGluR2-GFP, led to clear surface labeling with BG-Alexa-546 (Fig 4B; S4A). Co-expression of NB-SNAP, without the addition of the ss, was unable to label surface GFP-mGluR2 (**Fig S4B**). We next tested the ability of genetically-encoded, co-expressed NB-SNAP to optically control GFP-mGluR2. Similar to what was observed with purified NB-SNAP, BGAG_12_ produced clear photocurrents that were ∼20% in amplitude relative to saturating 1 mM glutamate (Fig 4C). Together these data demonstrate that anti-GFP NBs may be genetically encoded to efficiently target proteins for optical control. In addition to the advantage of permitting cell-type specific optical control in complex systems, this approach also provides the flexibility to either label a nanobody-GFP complex for optical control on the cell surface (using a membrane-impermeable photoswitch or fluorophore) or inside of a cell (using a membrane-permeable photoswitch or fluorophore) to probe the distinct roles and regulation of a signaling protein in different cellular locations, an increasingly appreciated aspect of GPCR function.[21, 22]

**Fig 4.**
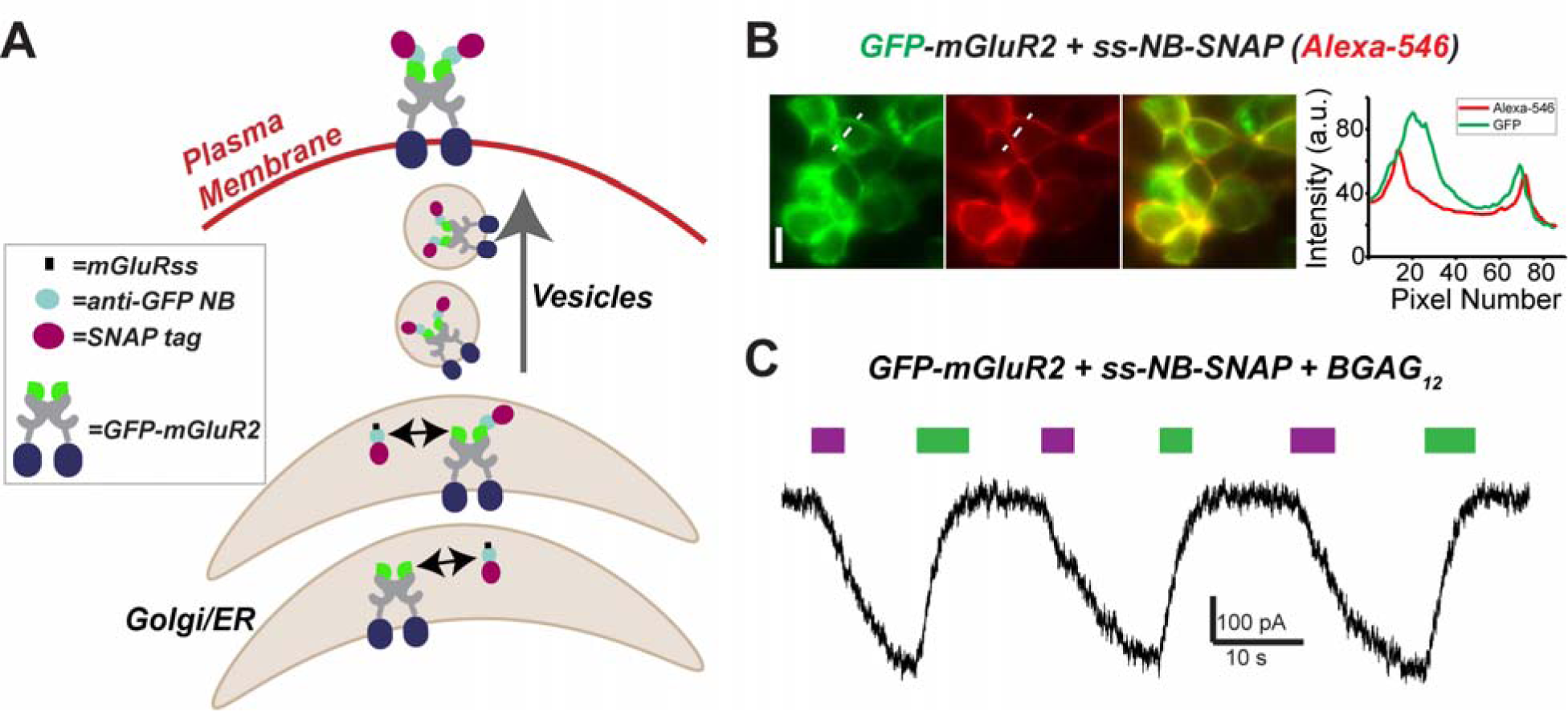
Genetic encoding of SNAP-tagged anti-GFP nanobodies allows targeting of extracellular GFP-tags for optical control. (A) Schematic showing co-trafficking and assembly of ss-NB-SNAP with GFP-mGluR2. (B) Representative images showing labeling of surface-expressed GFP-mGluR2 with co-expressed ss-NB-SNAP following labeling with Alexa-546. (C) Representative traces showing photoactivation of GFP-mGluR2 by BGAG_12_ conjugated to co-expressed ss-NB-SNAP.

## Discussion

In this study, we have established a new, general approach to targeting signaling proteins for optical control using nanobody-photoswitch conjugates (NPCs). In contrast to a recent study using an antibody to deliver an irreversible photosensitizer to inactivate AMPA receptors[23] this work establishes the suitability of immunochemistry for targeting a membrane protein for *reversible* optical control, opening the door to the manipulation of native proteins with high spatiotemporal precision. Similarly, Scholler et al[24] recently reported *allosteric* nanobodies to manipulate mGluR2, but these tools also lack reversibility and were not shown to permit genetic encoding for cell-type targeting, limiting their utility for precise, dynamic manipulation of receptors. In addition, the ability of NPCs to manipulate signaling via the *orthosteric* binding site via PORTLs permits a more physiological receptor perturbation through a defined, native mechanism.

Recent work has supported the idea that antibody-based approaches to manipulating GPCRs show great promise for treatment of a wide range of diseases, including various cancers and neurological disorders[25]. Importantly, the principles reported in this manuscript should be widely applicable *via* other types of antibodies and other receptor subtypes, and may ultimately be useful clinically by adding reversible spatiotemporal control to traditional antibody-drug conjugates (ADCs)[26]. Crucially, we demonstrate that NPCs can either be applied as purified complexes, which are most applicable in this clinical context, or as genetically-encoded tools that can be used to target specific cell types for maximal precision in mechanistic biological studies. While the ability to target NBs directly to wild-type, endogenous proteins of interest offers great potential, the targeting of proteins with incorporated GFP tags also offers many advantages. Strong antibodies or nanobodies with sufficient strength and specificity are not available for many antigens but GFP, for which exceptionally high affinity nanobodies have been developed, extensively characterized and applied *in vitro* and in living cells [13, 27] [28] has been successfully incorporated into many proteins, often in multiple different positions, that have been well-characterized functionally[29]. In addition, recent genetic advances based on the CRISPR-Cas9 system have made it increasingly feasible to incorporate GFP or other tags into native proteins either at the transgenic level or even in post-mitotic cells in the nervous system[30] [31]. Ultimately, the existing toolbox of photoswitchable ligands for SNAP and CLIP with different spectral properties, NBs and single chain antibodies for ubiquitous tags, along with the complement of genetic lines for targeting NB expression, should allow for sophisticated multiplexed experiments that can manipulate native signaling proteins to probe their roles in physiology with optimal precision.

## Materials and Methods

### Chemical Synthesis

Chemical synthesis was performed as previously described[8, 10].

### Cloning, Purification and Characterization of Nanobodies

All fusion proteins were cloned using standard cloning techniques. SNAP NB-fusions with an *N*-terminal Strep-tag and 10xHis-tag were cloned into a pBAD expression vector for bacterial expression, wtGFP with an *N*-terminal Strep-tag and a *C*-terminal 10xHis-tag was cloned into pET51b(+) for bacterial expression, and Complete amino acid sequences for constructs used can be found in the supplementary material. ss-NB-SNAP contains the mGluR5 signal sequence (“MVLLLILSVLLLKEDVRGSA”) at the *N*-terminus. For purification, SNAP-fused NBs were expressed in the *E. coli* strain DH10B and wtGFP was expressed in BL21(DE3) pLysS cell. All LB media contained ampicillin (100 µg/mL) for protein expression. A culture was grown at 37 °C until an OD 600 of 0.6 was reached at which point cells were induced with *L*-arabinose (0.02%, (w/v)) or IPTG (1 mM). Protein constructs were expressed overnight at 25 °C. Cells were harvested by centrifugation and sonicated to produce cell lysates. The lysate was cleared by centrifugation and purified by Ni-NTA resin (Thermofisher) and Strep-Tactin II resin (IBA) according to the manufacturer’s protocols. Purified protein samples were stored in 50 mM Hepes, 50 mM NaCl (pH 7.3) and either flash frozen and stored at −80 °C or stored at 4 °C. SNAP-tag was labeled with fluorescent dyes or photoswitchable compounds at 2-10 µM protein with a 2-5 fold molar ratio of fluorophores in 50 mM Hepes, 50 mM NaCl (pH 7.3). For *in vitro* experiments, excess label was removed by centrifugal filter devices with 10 KDa cutoff (Microcon YM-10, Millipore), and the labeled protein was stored in 50 mM Hepes, 50 mM NaCl (pH 7.3) at 4 °C.

Activity testing of purified nanobody-SNAP fusions against wtGFP was performed in 50 mM Hepes, 50 mM NaCl (pH 7.3) 0.05 mg/mL BSA, 0.05% Triton X-100 in black 384-well plates (Corning, 3820). Fluorescent intensities were recorded on a Spark 20M (Tecan) with excitation at 480 nm, emission at 535 nm, and bandwidths of 5 nm for both excitation and emission. The dose-response was fit with Eq.1.

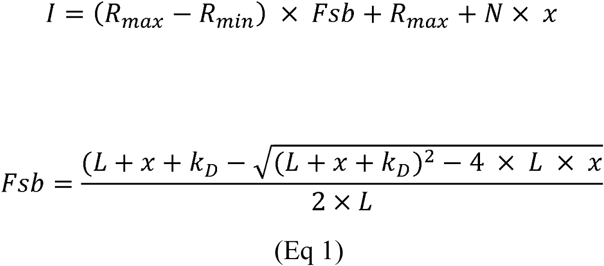

Where *I* is the Intensity of light at 535 nm, *R_max_* is the maximum light emitted at 535 nm, *R_min_* is the minimum light emitted at 535 nm, *L* is the concentration of wtGFP and fixed to 4 nM, *x* is the concentration of the nanobody-SNAP fusion protein, and *K_D_* is the calculated dissociation constant, and *N* accounts for non-specific binding. The fit was performed in Prism (GraphPad).

Competitive labeling and in-gel imaging was used to assay the efficiency of BGAG_12_ labeling of SNAP-tag NB-fusions. SNAP-NB or NB-SNAP (5 µM) were treated with BGAG_12_ (15 µM, 3-fold equivalence) in 50 mM Hepes, 50 mM NaCl (pH 7.3). Aliquots were taken at indicated time points and labeled with BG-TMR (150 µM, 30 equivalents) for 1 h. The reaction was then quenched with SDS-loading buffer, heated to 95 °C for 10 minutes, and stored at −20 °C until separated by SDS-PAGE. In-gel fluorescence of BG-TMR was measured on a ChemiDoc MP imager (Bio Rad) equipped with a CCD camera. Imaging was performed using green epifluorescent light and emission filters for Cy3 (605/50 nm.). The gel was then labeled with coomasie. Image analysis was performed with ImageJ (W. Rasband, NIH). Fluorescence Intensities in the Cy3 channel were background subtracted and normalized to the fluorescence with no BGAG_12_ present at time 0.

### Mass analysis

A Bruker maXis II ETD quadrupole-time-of-flight (QTOF) instrument with reverse phase liquid chromatography was used to measure accurate protein masses. All measurements were performed by injecting 5 µL of protein in 50 m M Hepes, 50 mM NaCl (pH 7.3) at a concentration of 5 µM. Proteins were first passed over a C8 wide pore column (100 mm) heated to 50 °C in a gradient of H_2_O + 1% formic acid (A)/MeCN + 1% formic acid (B) from 10 to 98% B over 6 minutes. The charge envelope of the protein within the corresponding spectrum was deconvoluted using Bruker’s DataAnalysis software with the Maximum Entropy deconvolution option enabled.

### Molecular Biology, Cell Culture, and Transfection

GFP-mGluR2 in pcDNA3 was produced using standard PCR-based techniques. In brief, enhanced GFP (eGFP) and a 17 amino acid glycine-rich linker were added after the mGluR2 signal sequence. HEK293T cells were maintained in DMEM media supplemented with 5% fetal bovine serum. Cells were seeded on 18 mm coverslips and transfected 12 hours later using Lipofectamine 3000 (Thermo Fisher). 0.3-0.7 µg each of GFP-mGluR2 DNA, GIRK1-F137S DNA (for electrophysiology experiments), and NB-SNAP (for NB co-expression experiments) was added to each well. Experiments were performed 24-48 hours after transfection.

### Fluorescence Imaging

Imaging was performed on an IX-73 microscope (Olympus) with a cellTIRF laser illumination system and an sCMOS ORCA-Flash4 v3.0 camera (Hamamatsu). GFP was excited using a 488 nm laser diode and a 561 nm DPSS laser was used for Alexa-546. Cellular imaging was performed in wide field mode using a 60x objective (NA=1.49). Data was acquired using cellSens software (Olympus) and analyzed using ImageJ. Prior to imaging, cells were incubated with purified, Alexa-546 labeled NBs (50 nM) for 45 minutes at 37 °C. For genetically-encoded ss-NB-SNAP experiments, BG-Alexa-546 was applied at 1 µM for 45 minutes at 37 °C. Single molecule pulldown of mGluR2 was performed as previously described[16]. Briefly, following immobilization using an anti-mGluR2 antibody images were acquired with a 100x objective (NA=1.49) at 20 Hz in TIRF mode and bleaching steps for individual molecules were manually analyzed using a previously described custom software [32].

### Patch Clamp Electrophysiology

Whole cell patch clamp electrophysiology was performed as previously described [10]. In brief, HEK 293T cells co-expressing GFP-mGluR2 along with GIRK1-F137S (homotetramerization mutant) were voltage clamped at −60 mV using an Axopatch 200B amplifier (Molecular Devices) attached to a 3-6 MΩ pipette filled with intracellular solution containing 140 mM KCl, 10 mM Hepes, 5 mM EGTA, and 3 mM MgCl_2_ (pH 7.4). Recordings were performed in high potassium extracellular solution containing 120 mM KCl, 25 mM NaCl, 1 mM MgCl_2_, 2 mM CaCl_2_, and 10 mM Hepes (pH 7.4). Cells were incubated with purified NBs (100 nM) and/or BGAG variants (10 µM) for 45 minutes at 37 °C. Prior to any recordings, cells were extensively washed with high potassium extracellular solution using a gravity-driven perfusion system, which was also used to control glutamate application. Light stimulation (∼1 mW/mm^2^) was provided by a coolLED pE-4000 illumination system that was controlled *via* pClamp software *via* a Digidata 1550B digitizer. Data were analyzed with Clampfit (Molecular Devices) and Prism (GraphPad).

## Acknowledgements

We thank Ehud Isacoff (UC Berkeley), Jeremy Dittman (Weill Cornell) and Philipp Leippe (LMU) for helpful discussion. We also thank Sebastian Fabritz (MPImR) for assistance with mass spectrometry. D.T. is grateful to the Center for Integrated Protein Science, Munich, and the European Research Council (ERC Advanced Grant 268795) for financial support. K.J. is grateful to finding from the Swiss Science Foundation, the NCCR Chemical Biology and EPFL. This work was supported by an R35 grant to J.L. from the National Institute of General Medicine (1 R35 GM124731). J.B. acknowledges support from the ‘EPFL Fellows’ fellowship programme co-funded by Marie Skłodowska-Curie, Horizon 2020 Grant agreement no. 665667.

## References

1. Tischer D, Weiner OD. Illuminating cell signalling with optogenetic tools. Nat Rev Mol Cell Biol. 2014;15(8):551–8. doi: 10.1038/nrm3837. PubMed PMID: 25027655; PubMed Central PMCID: PMCPMC4145075.

2. Deisseroth K. Optogenetics: 10 years of microbial opsins in neuroscience. Nat Neurosci. 2015;18(9):1213–25. doi: 10.1038/nn.4091. PubMed PMID: 26308982; PubMed Central PMCID: PMCPMC4790845.

3. Spangler SM, Bruchas MR. Optogenetic approaches for dissecting neuromodulation and GPCR signaling in neural circuits. Curr Opin Pharmacol. 2017;32:56–70. doi: 10.1016/j.coph.2016.11.001. PubMed PMID: 27875804; PubMed Central PMCID: PMCPMC5395328.

4. Broichhagen J, Frank JA, Trauner D. A Roadmap to Success in Photopharmacology. Accounts of chemical research. 2015. doi: 10.1021/acs.accounts.5b00129. PubMed PMID: 26103428.

5. Broichhagen J, Trauner D. The in vivo chemistry of photochromic tethered ligands. Current Opinion in Chemical Biology. 2014;21:121–7.

6. Reiner A, Levitz J, Isacoff EY. Controlling ionotropic and metabotropic glutamate receptors with light: principles and potential. Curr Opin Pharmacol. 2015;20:135–43. doi: 10.1016/j.coph.2014.12.008. PubMed PMID: 25573450; PubMed Central PMCID: PMCPMC4318769.

7. Levitz J, Pantoja C, Gaub B, Janovjak H, Reiner A, Hoagland A, et al. Optical control of metabotropic glutamate receptors. Nat Neurosci. 2013;16(4):507–16. doi: 10.1038/nn.3346. PubMed PMID: 23455609; PubMed Central PMCID: PMCPMC3681425.

8. Broichhagen J, Damijonaitis A, Levitz J, Sokol KR, Leippe P, Konrad D, et al. Orthogonal Optical Control of a G Protein-Coupled Receptor with a SNAP-Tethered Photochromic Ligand. ACS Cent Sci. 2015;1(7):383–93. doi: 10.1021/acscentsci.5b00260. PubMed PMID: 27162996; PubMed Central PMCID: PMCPMC4827557.

9. Berry M, Holt A, Levitz J, Broichhagen J, Gaub B, Visel M, et al. Restoration of patterned vision with a photo-engineered G protein coupled receptor. Nat Commun.

10. Levitz J, Broichhagen J, Leippe P, Konrad D, Trauner D, Isacoff EY. Dual optical control and mechanistic insights into photoswitchable group II and III metabotropic glutamate receptors. Proc Natl Acad Sci U S A. 2017;114(17):E3546–E54. doi: 10.1073/pnas.1619652114. PubMed PMID: 28396447; PubMed Central PMCID: PMCPMC5410775.

11. Beghein E, Gettemans J. Nanobody Technology: A Versatile Toolkit for Microscopic Imaging, Protein-Protein Interaction Analysis, and Protein Function Exploration. Front Immunol. 2017;8:771. doi: 10.3389/fimmu.2017.00771. PubMed PMID: 28725224; PubMed Central PMCID: PMCPMC5495861.

12. Manglik A, Kobilka BK, Steyaert J. Nanobodies to Study G Protein-Coupled Receptor Structure and Function. Annu Rev Pharmacol Toxicol. 2017;57:19–37. doi: 10.1146/annurev-pharmtox-010716-104710. PubMed PMID: 27959623; PubMed Central PMCID: PMCPMC5500200.

13. Kirchhofer A, Helma J, Schmidthals K, Frauer C, Cui S, Karcher A, et al. Modulation of protein properties in living cells using nanobodies. Nat Struct Mol Biol. 2010;17(1):133–8. doi: 10.1038/nsmb.1727. PubMed PMID: 20010839.

14. Mollwitz B, Brunk E, Schmitt S, Pojer F, Bannwarth M, Schiltz M, et al. Directed evolution of the suicide protein O(6)-alkylguanine-DNA alkyltransferase for increased reactivity results in an alkylated protein with exceptional stability. Biochemistry. 2012;51(5):986–94. doi: 10.1021/bi2016537. PubMed PMID: 22280500.

15. Keppler A, Gendreizig S, Gronemeyer T, Pick H, Vogel H, Johnsson K. A general method for the covalent labeling of fusion proteins with small molecules in vivo. Nat Biotechnol. 2003;21(1):86–9. doi: 10.1038/nbt765. PubMed PMID: 12469133.

16. Levitz J, Habrian C, Bharill S, Fu Z, Vafabakhsh R, Isacoff EY. Mechanism of Assembly and Cooperativity of Homomeric and Heteromeric Metabotropic Glutamate Receptors. Neuron. 2016;92(1):143–59. doi: 10.1016/j.neuron.2016.08.036. PubMed PMID: 27641494; PubMed Central PMCID: PMCPMC5053906.

17. Doumazane E, Scholler P, Zwier JM, Trinquet E, Rondard P, Pin JP. A new approach to analyze cell surface protein complexes reveals specific heterodimeric metabotropic glutamate receptors. FASEB J. 2011;25(1):66–77. doi: 10.1096/fj.10-163147. PubMed PMID: 20826542.

18. Kniazeff J, Bessis AS, Maurel D, Ansanay H, Prezeau L, Pin JP. Closed state of both binding domains of homodimeric mGlu receptors is required for full activity. Nat Struct Mol Biol. 2004;11(8):706–13. doi: 10.1038/nsmb794. PubMed PMID: 15235591.

19. Tang JC, Szikra T, Kozorovitskiy Y, Teixiera M, Sabatini BL, Roska B, et al. A nanobody-based system using fluorescent proteins as scaffolds for cell-specific gene manipulation. Cell. 2013;154(4):928–39. doi: 10.1016/j.cell.2013.07.021. PubMed PMID: 23953120; PubMed Central PMCID: PMCPMC4096992.

20. Irannejad R, Tomshine JC, Tomshine JR, Chevalier M, Mahoney JP, Steyaert J, et al. Conformational biosensors reveal GPCR signalling from endosomes. Nature. 2013;495(7442):534–8. doi: 10.1038/nature12000. PubMed PMID: 23515162; PubMed Central PMCID: PMCPMC3835555.

21. Irannejad R, Tsvetanova NG, Lobingier BT, von Zastrow M. Effects of endocytosis on receptor-mediated signaling. Curr Opin Cell Biol. 2015;35:137–43. doi: 10.1016/j.ceb.2015.05.005. PubMed PMID: 26057614; PubMed Central PMCID: PMCPMC4529812.

22. Thomsen AR, Plouffe B, Cahill TJ, 3rd, Shukla AK, Tarrasch JT, Dosey AM, et al. GPCR-G Protein-beta-Arrestin Super-Complex Mediates Sustained G Protein Signaling. Cell. 2016;166(4):907–19. doi: 10.1016/j.cell.2016.07.004. PubMed PMID: 27499021; PubMed Central PMCID: PMCPMC5418658.

23. Takemoto K, Iwanari H, Tada H, Suyama K, Sano A, Nagai T, et al. Optical inactivation of synaptic AMPA receptors erases fear memory. Nat Biotechnol. 2017;35(1):38–47. doi: 10.1038/nbt.3710. PubMed PMID: 27918547.

24. Scholler P, Nevoltris D, de Bundel D, Bossi S, Moreno-Delgado D, Rovira X, et al. Allosteric nanobodies uncover a role of hippocampal mGlu2 receptor homodimers in contextual fear consolidation. Nat Commun. 2017;8(1):1967. doi: 10.1038/s41467-017-01489-1. PubMed PMID: 29213077; PubMed Central PMCID: PMCPMC5719040.

25. Hutchings CJ, Koglin M, Olson WC, Marshall FH. Opportunities for therapeutic antibodies directed at G-protein-coupled receptors. Nat Rev Drug Discov. 2017;16(9):787–810. doi: 10.1038/nrd.2017.91. PubMed PMID: 28706220.

26. Beck A, Goetsch L, Dumontet C, Corvaia N. Strategies and challenges for the next generation of antibody-drug conjugates. Nat Rev Drug Discov. 2017;16(5):315–37. doi: 10.1038/nrd.2016.268. PubMed PMID: 28303026.

27. Kubala MH, Kovtun O, Alexandrov K, Collins BM. Structural and thermodynamic analysis of the GFP:GFP-nanobody complex. Protein Sci. 2010;19(12):2389–401. doi: 10.1002/pro.519. PubMed PMID: 20945358; PubMed Central PMCID: PMCPMC3009406.

28. Sommese RF, Hariadi RF, Kim K, Liu M, Tyska MJ, Sivaramakrishnan S. Patterning protein complexes on DNA nanostructures using a GFP nanobody. Protein Sci. 2016;25(11):2089–94. doi: 10.1002/pro.3020. PubMed PMID: 27538185; PubMed Central PMCID: PMCPMC5079250.

29. Giepmans BN, Adams SR, Ellisman MH, Tsien RY. The fluorescent toolbox for assessing protein location and function. Science. 2006;312(5771):217–24. doi: 10.1126/science.1124618. PubMed PMID: 16614209.

30. Mikuni T, Nishiyama J, Sun Y, Kamasawa N, Yasuda R. High-Throughput, High-Resolution Mapping of Protein Localization in Mammalian Brain by In Vivo Genome Editing. Cell. 2016;165(7):1803–17. doi: 10.1016/j.cell.2016.04.044. PubMed PMID: 27180908; PubMed Central PMCID: PMCPMC4912470.

31. Nishiyama J, Mikuni T, Yasuda R. Virus-Mediated Genome Editing via Homology-Directed Repair in Mitotic and Postmitotic Cells in Mammalian Brain. Neuron. 2017;96(4):755–68 e5. doi: 10.1016/j.neuron.2017.10.004. PubMed PMID: 29056297; PubMed Central PMCID: PMCPMC5691606.

32. Ulbrich MH, Isacoff EY. Subunit counting in membrane-bound proteins. Nat Methods. 2007;4(4):319–21. doi: 10.1038/nmeth1024. PubMed PMID: 17369835; PubMed Central PMCID: PMCPMC2744285.

